# Real-Time Flow Cytometry to assess qualitative and quantitative responses of oral pathobionts during exposure to antiseptics

**DOI:** 10.1101/2022.02.28.482436

**Authors:** I. Chatzigiannidou, J. Heyse, R. Props, P. Rubbens, W. Teughels, T. Van de Wiele, N. Boon

## Abstract

Antiseptics are widely used in oral healthcare to prevent or treat oral diseases, such as gingivitis and periodontitis. However, the incidence of bacteria being tolerant to standard antiseptics has sharply increased over the last few years. This stresses the urgency for surveillance against tolerant organisms, as well as the discovery of novel antimicrobials. Traditionally, susceptibility to antimicrobials is assessed by broth micro-dilution or disc diffusion assays, both of which are time-consuming, labor-intensive and provide limited information on the mode of action of the antimicrobials. The above-mentioned limitations highlight the need for the development of new methods to monitor and further understand antimicrobial susceptibility. Here, we used real-time flow cytometry, combined with membrane permeability staining, as a quick and sensitive technology to study the quantitative and qualitative response of two oral pathobionts to different concentrations of chlorhexidine, cetylpyridinium chloride or triclosan. Apart from the real-time monitoring of cell damage, we further applied a phenotypic fingerprint method to differentiate between the bacterial subpopulations that arose due to treatment. We quantified the pathobiont damage rate of different antiseptics at different concentrations within 15 minutes of exposure and identified the conditions under which the bacteria were most susceptible. Moreover, we detected species-specific and treatment-specific phenotypic subpopulations. This proves that real-time flow cytometry can provide information on the susceptibility of different microorganisms in a short time frame while differentiating between antiseptics and thus could be a valuable tool in the discovery of novel antimicrobial compounds while at the same time deciphering their mode of action.

## Introduction

Over the last decades the administration of antiseptics in oral healthcare for the treatment and prevention of oral diseases, such as gingivitis and periodontitis, has been intensified. However, there is increasing evidence that microorganisms become more tolerant to antiseptics, and this phenomenon is often combined with increased resistance towards antibiotics^1,2^. This stresses the need for, on the one hand, better surveillance for the antiseptic tolerance^3,4^ and, on the other hand, the acceleration of novel antimicrobial discovery.

Traditionally, antimicrobial susceptibility testing includes broth micro-dilution assays or disc diffusion assays^5,6^ with which the Minimum Inhibitory Concentration (MIC) is determined. These methods are confronted with limitations. They are based on the active growth of a bacterial strain under specific antiseptic concentrations and are therefore time-consuming as they need at least 24 hours of incubation from antiseptic application until endpoint measurement. Furthermore, they cannot differentiate between bacteriostatic and bactericidal conditions. To identify bactericidal conditions (determining the Minimum Bactericidal Concentration – MBC) another 24 hours are required. Additionally, these bulk based methods do not permit the detection of differences between bacterial subpopulations. Nonetheless, it is reported that even isogenic bacterial populations can harbor phenotypically variable subpopulations with different levels of tolerance towards antimicrobials^7,8^ and such differential response can affect the treatment outcome. Furthermore, the above-mentioned methods are not informative on the mode of action of tested compounds. Consequently, we need rapid antimicrobial susceptibility testing methods which can further provide information on the subpopulation level or the compounds mode of action. These methods can facilitate better infection management, antimicrobial tolerance surveillance and research for novel compounds.

Flow cytometry has already been described as a method to detect antimicrobial susceptibility^9,10^. One of the main advantages of this method is that it provides a large amount of quantitative data in a short time frame as it can measure hundreds to thousands of cells in a few seconds. Besides the rapid measurement, flow cytometry protocols can further accelerate testing because they do not require long incubation times in contrast to the conventional methods, allowing to move from 24-48 hours to 1-2 hours incubation window^10,11^. Moreover, with the use of appropriate dyes, it can directly detect cell damage and lysis thus enabling the study of bactericidal or bacterolytic activity^12^. In addition, the application of real-time flow cytometry could increase the time resolution as it enables the immediate observation of physiological changes of a given microbial subpopulation within a few seconds^13^.

In this study, we evaluated real-time flow cytometry as an accurate and fast method to study the response of two oral pathobionts when exposed to antiseptics commonly used in oral care. *Aggregatibacter actinomycetemcomitans* was exposed to chlorhexidine (CHX) and cetylpyridinium chloride (CPC), while *Streptococcus mutans* was tested under the above mentioned antiseptics and triclosan. By exposing the bacteria to the antiseptics for 15 minutes, we evaluated the quantitative and qualitative response in real-time under different concentrations of the antiseptics by respectively detecting cell damage and observing the dynamics of the phenotypical subpopulations that arose during the exposure to the antimicrobial.

## Results

To demonstrate the use of real-time flow cytometry as a method to assess the susceptibility of oral bacteria to antiseptics, two species with different physiological properties that are linked to oral diseases were incorporated in the study. More specifically, *Streptococcus mutans*, a Gram^+^ coccus linked to dental caries^14^, and *Aggregatibacter actinomycetemcomitans*, Gram^-^ coccobacillus that has been associated with more aggressive forms of periodontitis^15^, were used. The bacteria were subjected to different antiseptics, commonly found in oral care products, in different concentrations. A cell membrane permeability staining protocol was employed to distinguish between intact and damaged cells. SYBR Green I was used to stain all cells, while Propidium iodide, which only penetrates cells with disrupted cell membrane, was used to stain damaged cells. After staining, the cells were exposed to the respective antiseptics and continuously measured by flow cytometry for 15 min. Due to the antiseptic-induced cell damage, the cells were getting gradually stained with Propidium iodide, thus allowing the real-time observation of cell damage.

### Real-time determination of the cell damage rates

Our first objective was to measure the cell damage rate for each treatment, i.e. antiseptic type and concentration. We quantified the number of intact and damaged cells over time and segmented the 15 minutes continuous data in smaller time frames of 30 seconds. Two gates were drawn manually in the Green (FL1) vs. Red fluorescence intensity (FL3) density plots corresponding to the intact and damaged cell populations for each species (Supplementary Figure 1). To define the gates, we used a non-treated sample as a control for the intact population and a heat-killed sample as a control for the damaged population. Subsequently these gates were used to count the number of intact and damaged cells in each 30 seconds-frame.

The number of intact cells was used to further calculate the cell damage rate. More specifically, we expressed the data as the percentage of surviving cells which corresponds to the ratio of intact cells at each time point over the intact cells from the non-treated control samples (Figure 1 and 2). To define the rate of cell damage of each treatment, log-logistic models were fitted on the surviving cell percentage data. The Hill coefficient (slope) and the effective time 50 (ET50), calculated based on the model, were used to evaluate the effect of each treatment on the survival of the bacterial cells. The Hill coefficient describes the steepness of the curve, while the ET50 indicates the time after which 50% of the cells were damaged. A higher Hill coefficient means that the slope is steeper which corresponds to a faster response. A smaller ET50 value means that less time was needed to get half of the bacterial population in a damaged state.

**Figure 1:**
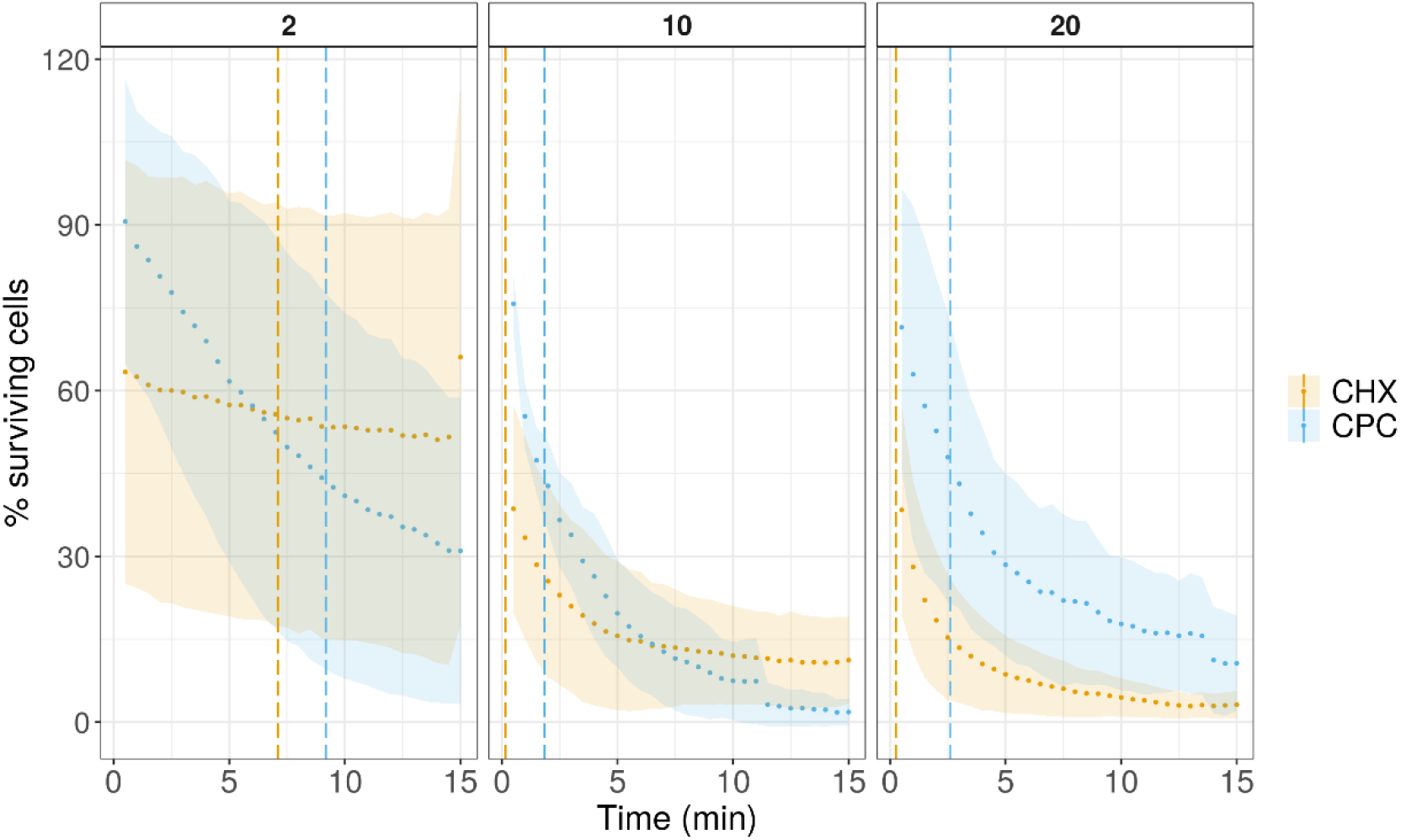
The percentage of surviving cellsl (intact cells) of *A. actinomycetemcomitans* over the time of 15 min for two different antiseptics, chlorhexidine (CHX) and cetylpyridinium chloride (CPC) in three different concentrations (2 mg/mL, 10 mg/mL and 20 mg/mL). The points represent the average of three replicates, while the ribbon the sample standard deviation. The vertical lines represent the ET50 time point.

**Figure 2:**
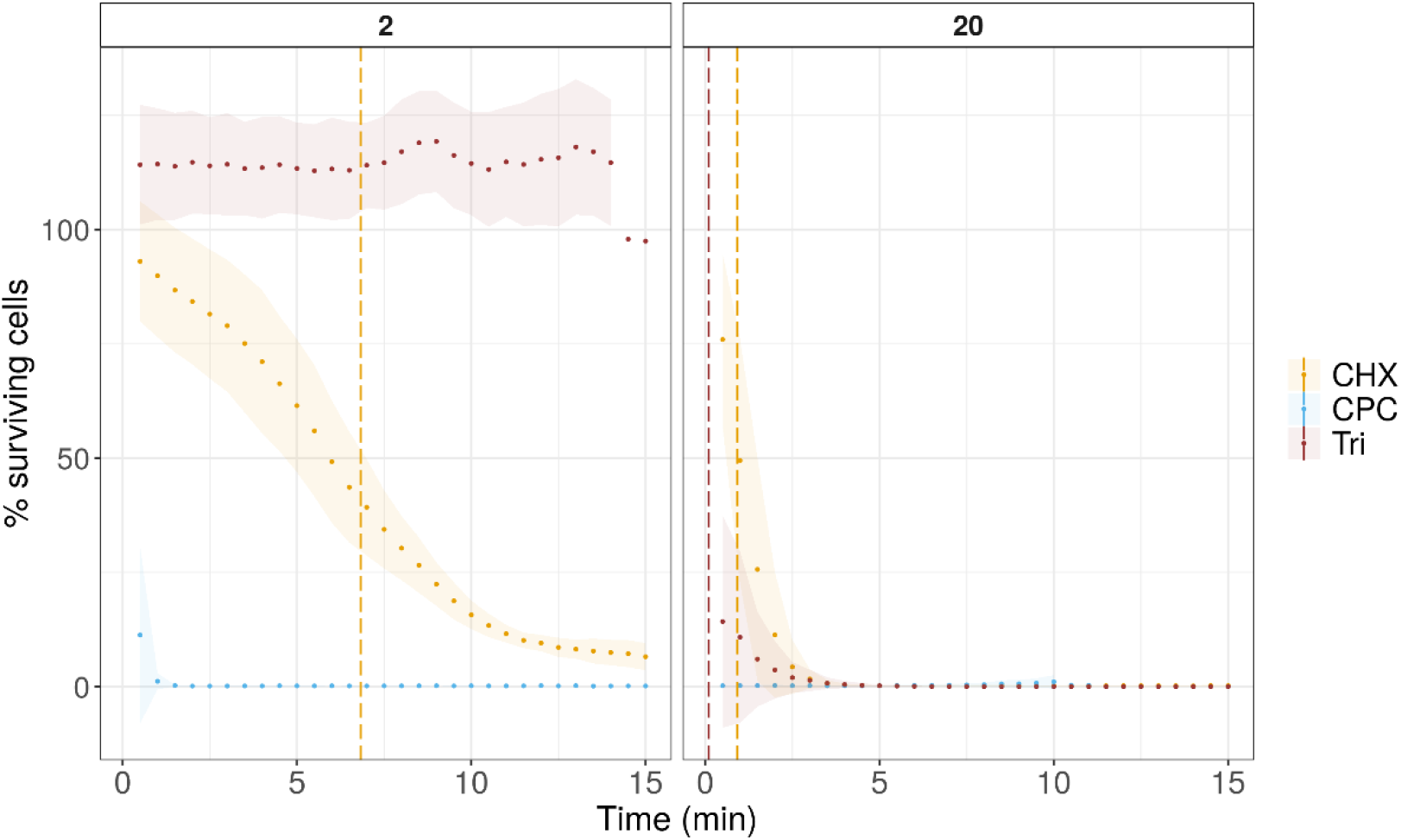
The percentage of surviving cells (intact cells) of *S. mutans* over the time of 15 min for two different antiseptics, chlorhexidine (CHX), cetylpyridinium chloride (CPC) and triclosan in two different concentrations (2 mg/mL and 20 mg/mL). The points represent the average of three replicates, while the ribbon the sample standard deviation. The vertical lines represent the ET50 time point.

Although CHX caused faster cell damage on *A. actinomycetemcomitans* cells, CPC could cause more damage over time when 2 and 10 mg/mL were applied. After choosing a three-parameter log-logistic model as the best-fitted model (Supplementary Figure 3A) and based on the slope steepness, we observed that CHX treatment, independent of concentration, exhibited a lower Hill coefficient, less steep killing curve, than the CPC treatment. The Hill coefficient for CHX was increasing with increasing concentrations, while the opposite was true for CPC (Figure 3). On the other hand, all CHX treatments exhibited a lower ET50 than the equal concentrations of CPC (Figure 3). Besides calculating the time-response curves for different treatments, the dose-response was calculated for samples that were exposed to treatment for 10 minutes. This time was chosen to ensure sufficient time from the exposure had passed. A three-parameter log-logistic model was used to calculate the Hill coefficient and the ED50, the dose for which 50% of cell damage should be observed (Supplementary Figure 3B). CPC had an ED50 of 1.5 mg/mL, while CHX had an ED50 of 1.9 mg/mL (Table 1), indicating that a slightly smaller concentration of CPC is needed to reach 50% of cell damage after 10 min of treatment.

**Figure 3:**
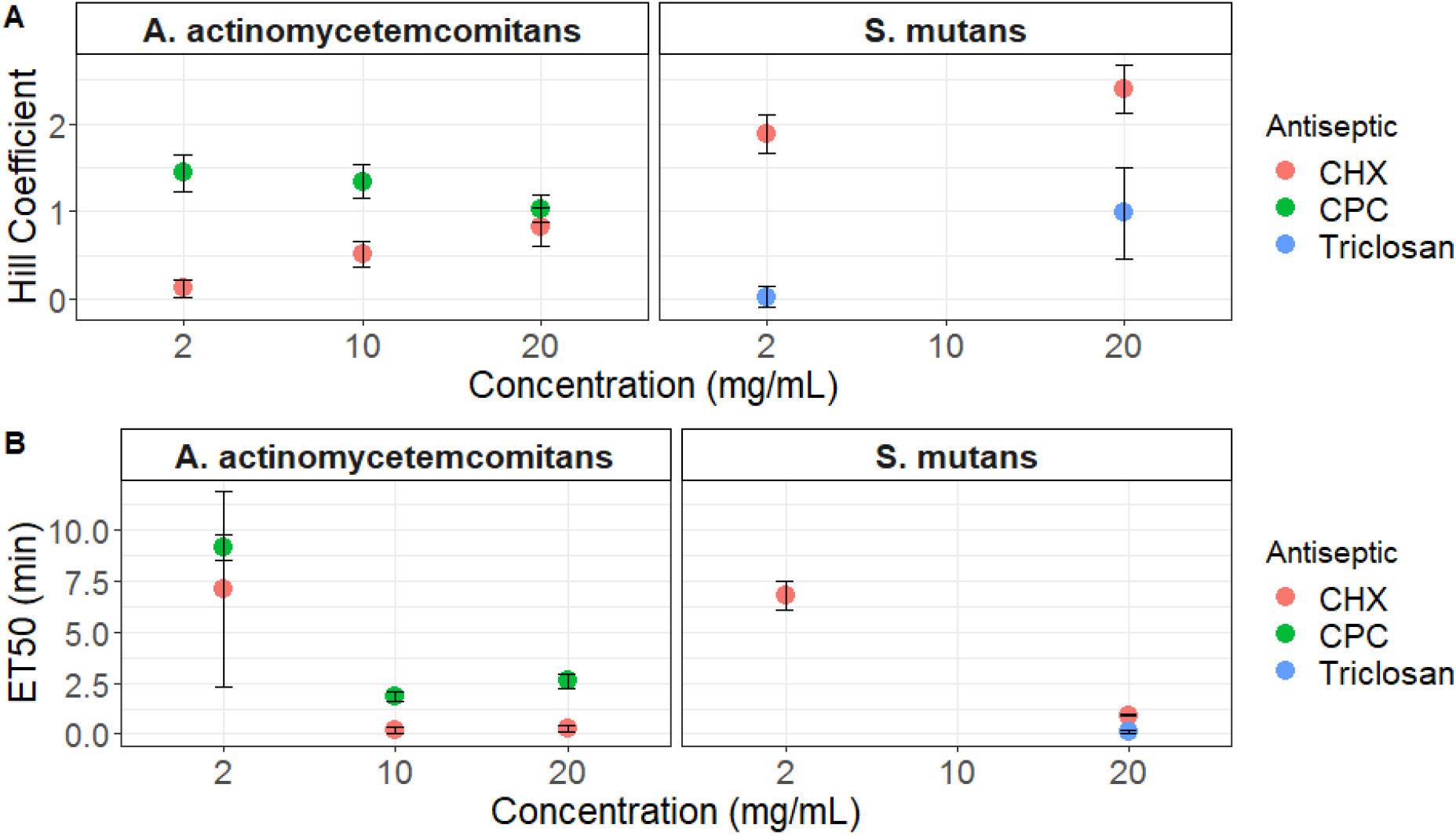
A. The Hill coefficient (slope of the curve) and B. Median Effective Time (ET50) as they were calculated based on time-dependent log-logistic models that were fitted in the cell damage curves of *A. actinomycetemcomitans* and *S. mutans* for each treatment. The values for *S. mutans* exposed to CPC could not be calculated as all cells were damaged in the beginning of the measurement. The values for the same species exposed to 2 mg/mL triclosan are not depicted as no killing was observed. The points represent the mean value of three biological replicates, while the error bars the standard deviation.

**Table 1:** The mean and standard deviation of the Hill coefficient (slope of the curve) and Median Effective Dose (ED50) as they were calculated based on dose-dependent log-logistic models that were fitted in the cell damage curves of *A. actinomycetemcomitans*.

| Species | Antiseptic | Time (min) | Hill coefficient | ED50 (mg/mL) |
| --- | --- | --- | --- | --- |
| *A. actinomycescomitans* | CHX | 10 | 1.29 (0.71) | 1.9 (0.73) |
| *A. actinomycescomitans* | CPC | 10 | 0.83 (0.5) | 1.52 (1.06) |

Different patterns were observed when *S. mutans* was exposed to the same antiseptics. Besides, this strain was also exposed to triclosan, which has a different mode of action than CHX and CPC. Exposure to CPC, either 2 or 20 mg/mL lead to immediate damage of the majority of the cells (< 99%) (Figure 3.2). As a consequence, we could not fit a time-response curve on this data and these conditions were not used in the next steps of the analysis. A four-parameter log-logistic model was used to calculate the rate of cell damage under treatment with the other two antiseptics (Supplementary Figure 8.4C). A higher Hill coefficient was observed for 20 mg/mL CHX compared to 2 mg/mL (Figure 3.3). The Hill coefficient of 2 mg/mL triclosan was almost 0, (0.02 ± 0.12), because no cell damage occurred during this treatment. Concerning the dose-dependent data, a good model that would properly describe the phenomenon could not be fitted, probably because of the lack of a range of concentrations that will capture different phases of killing.

Another gate representing the total bacterial cells, thus separating both intact and damaged populations from the background, was used to evaluate whether the tested conditions lead only to cell damage or also to cell lysis. For most of the tested conditions, minimal cell lysis was observed (< 10%) (Supplementary Figure 8.5 & 8.6). However, 20 mg/mL triclosan rapidly caused cell lysis of the *S. mutans* cells (Supplementary Figure 8.6).

### Flow-cytometric phenotypic fingerprinting

In the previous section, real-time flow cytometry data was used to calculate the effectiveness and rate of permeabilization of different antiseptic treatments by dividing the cell populationin intact and damaged subpopulations. Nevertheless, we noticed that the response to the different antiseptics was much more dynamic than could be captured by this binary classification and that intermediate physiological phenotypes appeared through time. To consider this information in our analysis, we employed an alternative approach which is based on the phenotypic fingerprint of the samples. Instead of the binary split in the manually designed gates, a Gaussian Mixture model was used to identify different physiological subpopulations (phenotypes).

After denoising the data based on the ‘total bacteria’ gate (Supplementary Figure 2), a representative subset of samples was chosen to estimate the parameters for the Gaussian Mixture Model. The information of four parameters, i) forward scatter (FSC), ii) side scatter (SSC), iii) green fluorescence intensity (FL1) and iv) red fluorescence intensity (FL3), was used and the model was set to identify 20 phenotypes (i.e. Gaussian mixtures). Including the forward and side scatter measurements on the model provided an extra layer of information about morphological differences between the subpopulations. By observing the mean value of each phenotype in the four channels, we found that separation was based on both fluorescence intensity and scatter information (Figure 4). Subsequently, the model was used to calculate the abundance of the different phenotypes in all time points.

**Figure 4:**
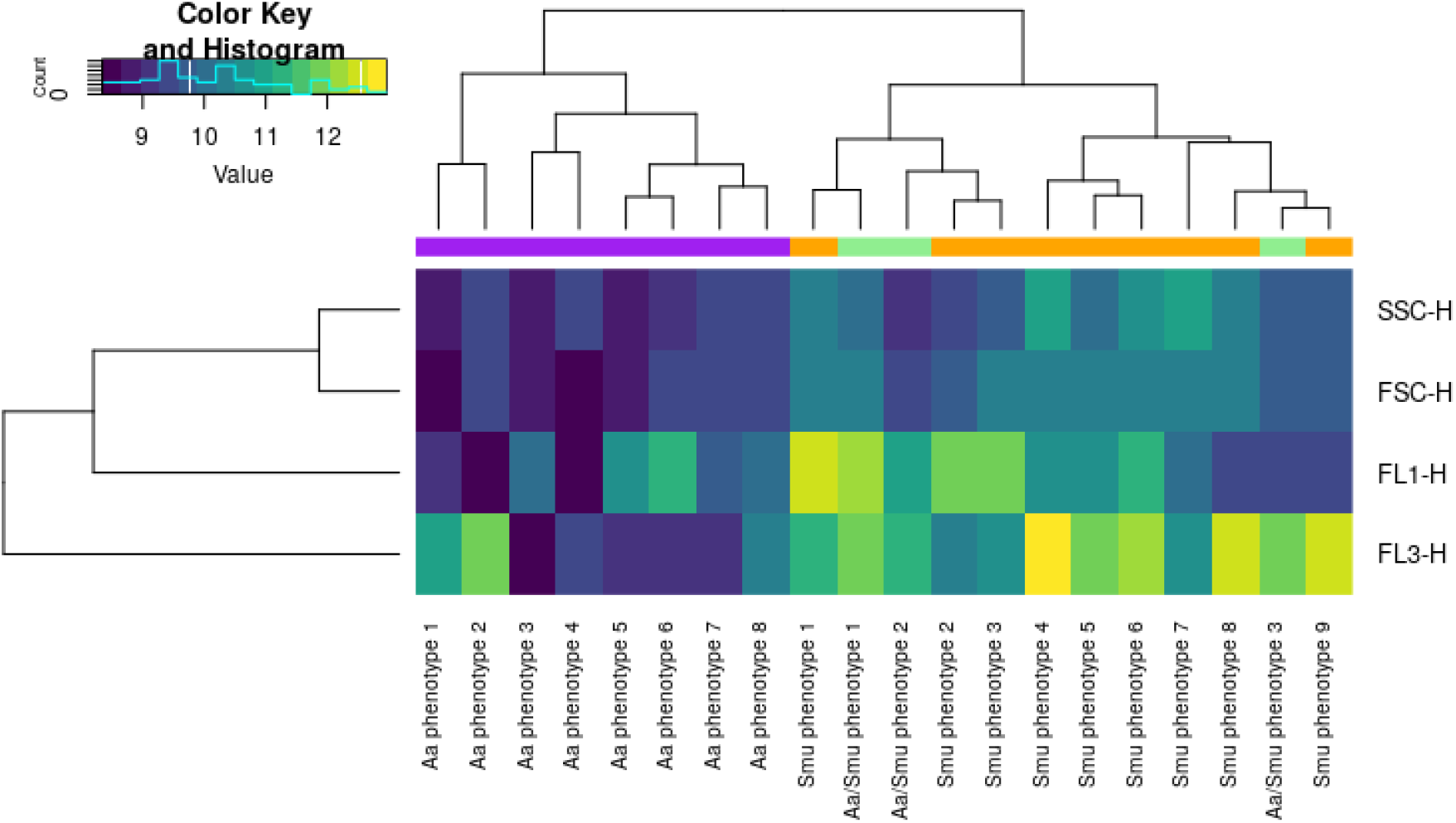
Heat map representing the mean values (a.u.) for each parameter (FL1, FL3, FSC, SSC) for the 20 phenotypes as they have been predicted by the Gaussian mixture model, when allowed to identify a maximum of 20 phenotypes using 1000 cells per sample as a training set. Phenotypes with higher red fluorescence represent damaged cells, while phenotypes with high green but low red fluorescence represent intact phenotypes. The rest of the phenotypes represent intermediate states most of which appeared during treatment. The color panel indicate whether the phenotypes were more abundant in *A. actinomycemcomitants* samples (purple) or in *S. mutans* samples (orange) or in both (green) and the phenotypes have been named accordingly.

The phenotypes clustered in two main groups according to the mean values of the four parameters (Figure 4) and the members of the one group (names Aa phenotype 1-8) were more abundant in the *A. actinomycetemcomitans* samples, while the members of the other group were more abundant in the *S. mutans* samples (named Smu phenotype 1-9), except from three phenotypes that were abundant in samples of both species (named Aa/Smu phenotype 1-3). Aa phenotypes 3, 5 and 6 were the ones describing the intact *A. actinomycetemcomitans* under no stress. We observed a transient shift from these phenotypes with the higher green fluorescence to the newly appearing phenotypes, when *A. actinomycetemcomitans* was exposed to the antiseptics. In the samples treated with CHX, these phenotypes had similar values of green and red fluorescence (Aa/Smu phenotype 1 and 2). The shift to these newly appearing phenotypes was slower or faster depending on the antiseptic concentration (Figure 5). We also observed that Aa phenotypes 7 and 8 appeared only under 10 and 20 mg/mL CPC and in a very low abundance under 2 mg/mL but were not present under treatment with CHX. Finally, it is noteworthy that the heat-killed cells clustered in two completely different phenotypes (Aa phenotype 1 and Aa/Smu phenotype 3) and not together with the cells damaged by antiseptics.

**Figure 5:**
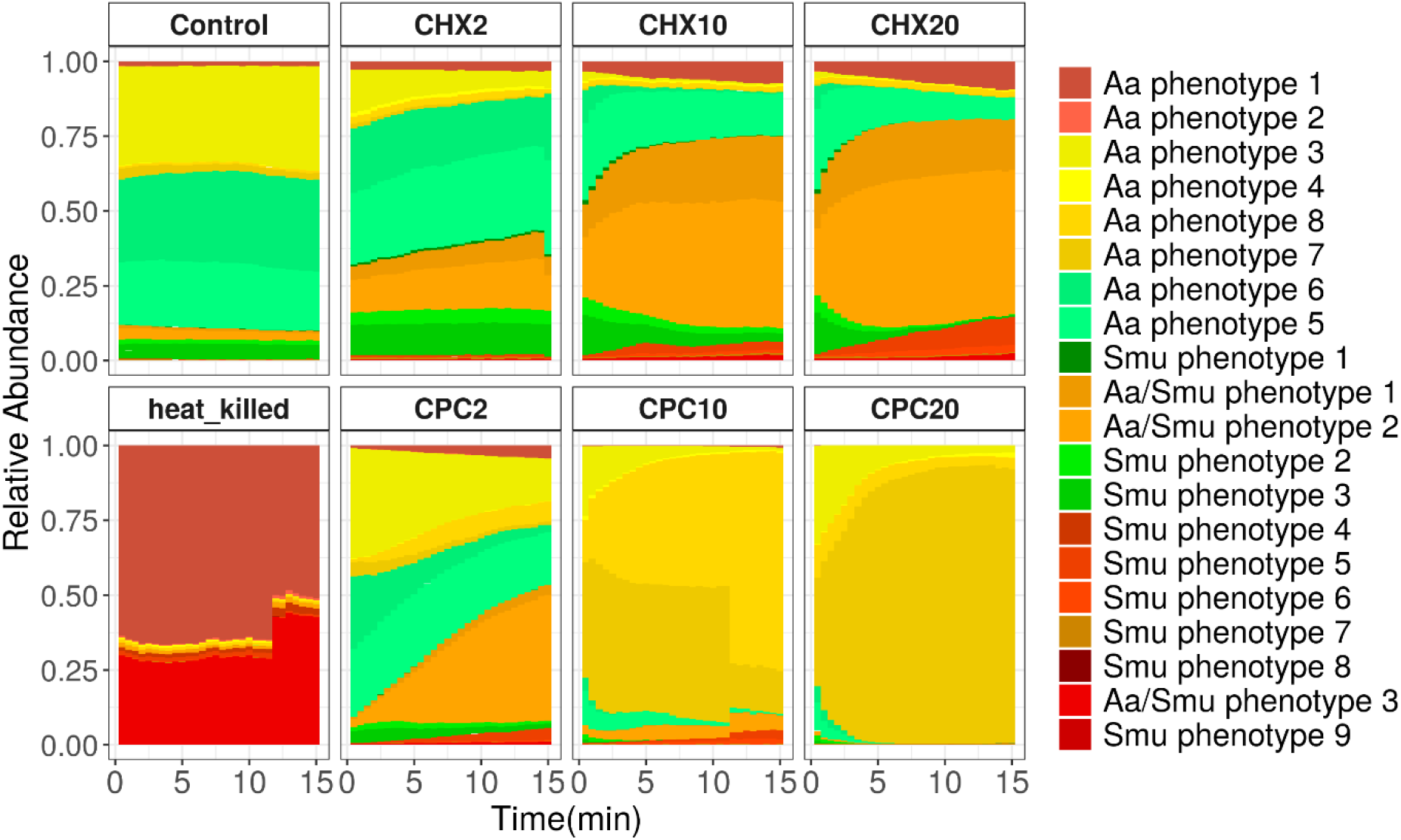
The relative abundances of phenotypic subpopulations in the different treatments and concentrations of *A. actinomycetemcomitans* as they have been estimated based on the Gaussian mixture model, which was allowed to identify a maximum of 20 phenotypes and trained in a subsection of the data using 1000 cells per sample. The bars represent the average of three biological replicates. The information about the mean value of the two fluorescence parameters was used to accordingly color the phenotypes. Samples with high red fluorescence are in different shades of red. Samples with high green and low red fluorescence are in shades of green. Samples with low green and red fluorescence are in shades of yellow, whille samples of median green and red fluorescence in shades of orange.

Intact non-stressed *S. mutans* cells were clustered in different phenotypes with the most abundant being Smu phenotypes 1,2 and 3. When *S. mutans* was exposed to CHX, different phenotypes appeared over time, moving from ones characterized from higher green fluorescence to ones with higher red fluorescence (Figure 6), while their abundances were dependent on the antiseptic concentration. Interestingly a range of different phenotypes was also observed in the conditions that all cells were clustered as damaged according to the previous method (Supplementary Figure 7). This was more clear for the CPC-treated *S. mutans* cells were all damaged from the first time point. However, differences between the damaged cells were observed with this method (Figure 6), with, for example, Smu phenotype 5 to be the most abundant under treatment with 20 mg/mL CPC and to be mainly found in this treatment. In addition, we observed that Smu phenotypes 5 and 6 that mostly characterized 20 mg/mL CPC treatment clustered together with Smu phenotype 4, more abundant in 20 mg/mL CHX, based on their mean values (Figure 3). The latter probably indicates that high concentrations of the two antiseptics induce similar but distinct cell subpopulations.

**Figure 6:**
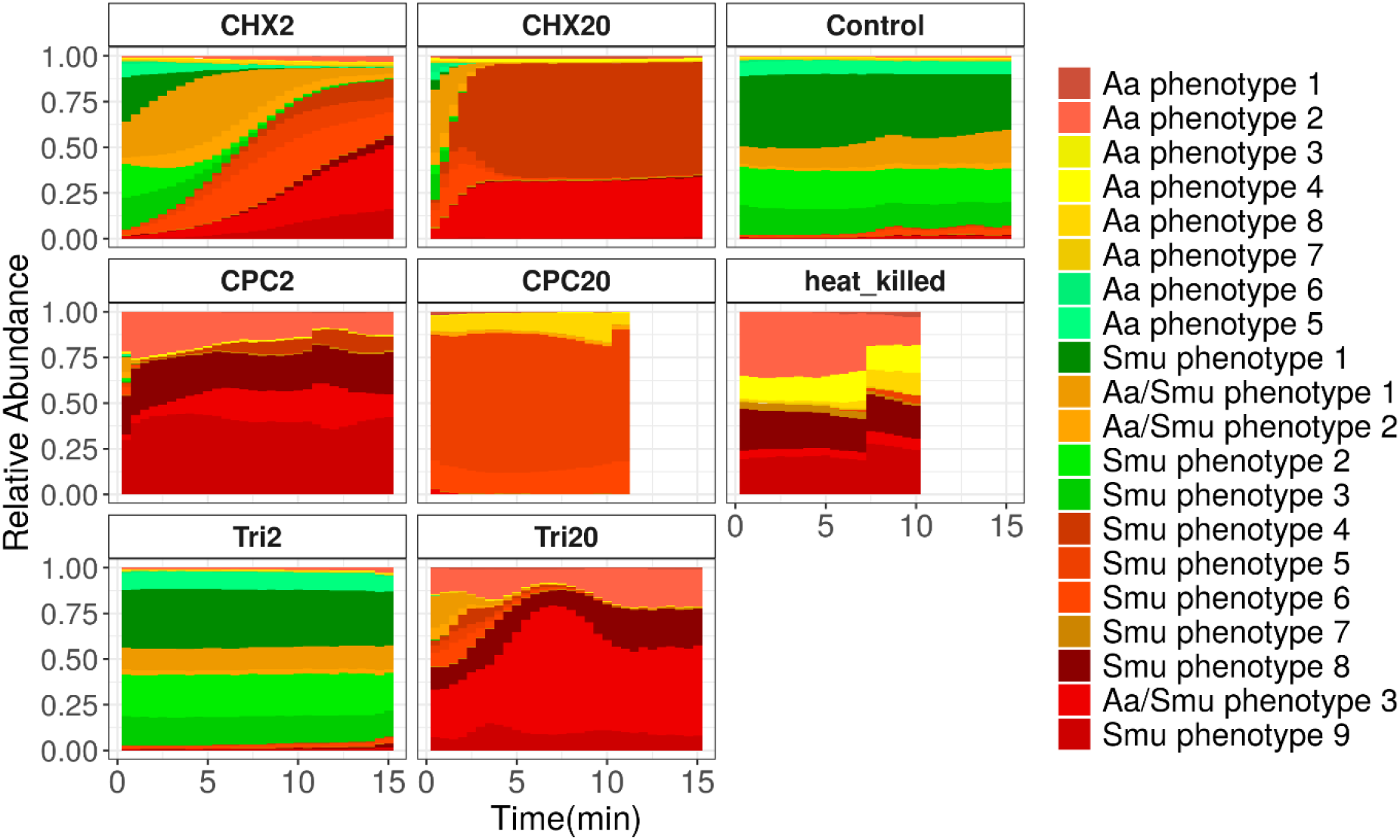
The relative abundances of phenotypic subpopulations in the different treatments and concentrations of *S. mutans* as they have been estimated based on the Gaussian mixture model, which was trained in a subsection of the data using 1000 cells per sample. The bars represent the average of three biological replicates. The information about the mean value of the two fluorescence parameters was used to accordingly color the phenotypes. Samples with high red fluorescence are in different shades of red. Samples with high green and low red fluorescence are in shades of green. Samples with low green and red fluorescence are in shades of yellow, whille samples of median green and red fluorescence in shades of orange.

Hence, with cytometric fingerprinting, we could observe the dynamic shift from phenotypes that were describing intact cells to a range of damaged cells phenotypes. Most importantly, we detected phenotypes that were either compound-specific or concentration-specific.

## Discussion

Flow cytometry is a quick and reliable method for determining microbial susceptibility of clinically important bacterial isolates^9,12,16^, for detecting bactericidal conditions^12^ and for accelerating prognosis in a clinical setting from 1-2 days to a couple of hours^10,11^. The combination of viability dyes (Syto9/PI) ^12,17^ or membrane polarization dyes (DiOC_n_(3), DiBAC_4_(3))^9^ allows corroborating membrane permeabilization and pore formation under certain stress. Different methods are in place to quantify the antimicrobial induced changes, such as manual gating^12^ or averaging ratios of the green and red fluorescence intensity to calculate the percentage of damaged cells^17^. Yet, our current approach of combining dynamic flow cytometry analysis with SGPI staining allowed us to achieve unprecedented real-time monitoring of membrane permeabilization of the bacterial population of two oral species during exposure to the three different antiseptics.

Initially, the killing dynamics upon exposure to a certain antiseptic were calculated based on the number of intact cells through time. *A. actinomycetemcomitans* was exposed to two different antiseptics, CHX and CPC and it was observed that CHX was faster in damaging the bacterial cells as it was indicated by the ET50 values (Figure 3). However, the rate of damage was decreasing over time, as this was calculated by the Hill coefficient (Figure 3). Contrastingly, when the two antiseptics were evaluated based on the concentration needed to decrease the cell population by half, it was found that slightly less CPC was needed to achieve the same effect (Table 1). Our findings suggest that, although CHX treatment acted more rapidly than CPC in the beginning, CPC was slightly more potent than CHX.

*S. mutans* cells exhibited three different patterns of response depending on the supplemented antiseptic. Immediate cell permeabilization was observed with CPC, gradual permeabilization over 15 min with CHX and a biphasic response with triclosan, for which one concentration, namely 20 mg/mL, led to immediate permeabilization while 10x lower concentration, 2mg/mL, caused no cell damage. Triclosan at low concentrations acts by blocking lipid synthesis via inhibition of enoyl-acyl carrier protein (ACP) reductase (FabI) ^18^ but at high concentrations seems to act simultaneously on multiple targets, such as inhibition of lipid, RNA, protein synthesis and membrane perturbations^19^. A different mode of action between lower and higher concentrations could be the reason for the biphasic response we observed. It is also important to note that 20 mg/mL triclosan was the only treatment under which cell lysis was observed.

Despite the substantial information that was acquired by this approach, some limitations must be considered. Short inactivation times of less than 1 minute prevented us from fitting a mathematical model to the antimicrobial effects, thus making the approach non-applicable for high concentrations of antiseptics. In addition, the subpopulations of intact and damaged cells were based on the 2-dimensional space of green vs red fluorescence intensity and were designed manually. Nonetheless, manual gating is subjected to bias ^20^and antimicrobial stress can affect not only cell permeability but also cell morphology ^21^. Moreover, by splitting the cells into two subpopulations, intact or damaged, we enforced a discretization of the data, which in reality represented a continuous spectrum of phenotypes with intermediate levels of cell permeability. Hence, meaningful biological information was not taken into account with this approach.

To overcome the above limitations, i.e. avoid the bias of manual assigning populations and make use of more dimensions, a cytometric fingerprinting approach was subsequently applied to the data. Fingerprinting techniques have been successfully used in the past to discriminate between different bacterial strains or the same strain in different physiological conditions^22,23^. They are superior to manual gating because they are not subjected to user bias^20^ and, in addition, they allow to observe phenotypical changes that might not be captured by the use of a binary system for intact/damaged classification^20^. Moreover, they take advantage of the multi-parametric measurements using both fluorescence and scatter information. In the past, very few efforts have been made to combine more than two-dimensions in susceptibility studies by flow cytometry, such as the study by Huang et al, 2015 which use the three-dimensional data to calculates the change between control and treated conditions as an one-dimensional distance^16^.

Here we used a Gaussian Mixture Model approach, to partition our data into different phenotypic subpopulations, which allows for the use of a smaller number of phenotypes as compared to the previous approaches^24^ and thus can better reflect the complexity of the physiological changes and cell damage caused by the treatment. Four parameters were taken into account for identifying different phenotypes (forward and side scatter and green and red fluorescence intensity), which provided information on the morphology, the nucleic acid content and the membrane permeability of every single cell. Both fluorescence intensity and scatter played a role in the separation of the phenotypes. This is in accordance with previous studies that have shown that stress, which results in cell wall damage, can alter the morphology of a bacterial cell^21,25^.

Applying the Gaussian Mixture Model flow cytometric fingerprinting method increased the resolution of our data analysis and revealed further information on how each treatment affected the phenotypic fingerprint of the two bacterial populations. Our results indicate that the relative abundances of the different phenotypes were dependent on the i). bacterial species, ii) antiseptic and concentration of antiseptic used and iii) the time after the application of the treatment. This method also revealed how the cells pass through different phenotypical stages with different values of cell permeabilization and cell morphology, revealing the dynamic process of antimicrobial action (Supplementary Figure 5 and 6). We noticed that the two main groups, in which the phenotypes were clustered (Figure 4), could be linked to one or the other species, which suggests that phenotypic differences of the two species are even larger than their state after antiseptic treatment. This could be explained by the lower forward and scatter values of the phenotypes mainly abundant on *A. actinomycetemcomitans* samples, which suggests a smaller size of this species. Additionally, the distinct phenotypes between antiseptic-treated *A. actinomycetemcomitans* and *S. mutans* could be explained by the differential action of the antiseptics against Gram^+^ and Gram^-^ species, such as the location where the cell wall is disrupted ^26^. These findings indicate that the method could be applied for species identification together with susceptibility testing, which might be an advantage in the case of clinical isolates.

Some phenotypes only appeared under one of the antiseptic treatments and not another, such as Aa phenotypes 7 and 8 that appeared only under CPC treatment of *A. actinomycetemcomitans* and Smu phenotype 5 under CPC treatment of *S. mutans*. We hypothesize that these distinct phenotypes could be linked to differences in the mode of action of the compounds. CHX is a bisbiguanide compound and CPC is a quaternary ammonium compound. Both of the compounds are positively charged and their mode of action is linked to their ability to bind in the negatively charged cell membrane, destabilizing the membrane and creating pores^27^. The only difference between the two compounds is that the hydrophobic region of CPC becomes solubilized within the hydrophobic core of the cell membrane while the hydrophobic region of chlorhexidine does not^28^. Therefore no major differences in the damaged phenotype would be expected from the use of the two compounds. However, for both species, it was apparent that CHX treatment resulted in phenotypes with higher red fluorescence than the ones that appeared under CPC treatment (Figure 3). Different pore sizes induced by each antiseptic could lead to the incorporation of different amounts of Propidium Iodide in the cells and could potentially explain the distinct phenotypes. Concerning triclosan treated *S. mutans*, when damage was observed, 20 mg/mL triclosan, the most abundant phenotype was Aa/Smu phenotype 3, which was not so abundant under treatment with CPC and CHX. Nevertheless, the mode of action of triclosan in high concentrations is not clear. Generally, information on the exact mode of action is lacking for many antiseptics and the above-described pipeline could help in understanding the physiological changes that antiseptics induce in microbial cells. Furthermore, flow cytometric fingerprinting can be used to detect persister subpopulations, by distinguishing phenotypes whose cell concentrations do not change overtime. Identifying antimicrobial compounds that will specifically target persister subpopulations can be of major importance as the increased tolerance of those subpopulations to antimicrobials hinter the efficacy of antimicrobial chemotherapy^29^.

In general, in the present study, we demonstrated the applicability of real-time flow cytometry for the study of antiseptic susceptibility. By using two different approaches, our analysis provided information on the rate of cell damage under different concentrations of antiseptics and allowed the comparison between treatments, while at the same time permitted the observation of the physiological changes induced by the different compounds over 15 minutes of exposure. We propose that the potential of this method can be further strengthened with the use of alternative staining dyes, e.g. membrane polarity stains, that can detect different physiological changes. Additionally, this method can be applied for other environmental stressors with clinical or biotechnological relevance that induce phenotypic changes, e.g. osmotic stress, UV, temperature. The temporal resolution can prove extremely beneficial to observe and quantify the dynamics at high concentrations of the stressor where cell response is instantaneous and thus distinct time points are insufficient to capture the intermediate phenotypic states. We believe that our study may trigger the broader use of real-time flow cytometry in the study of antimicrobial susceptibility, as it could be applied for the detection of resistant strains or in the quest for novel antimicrobial compounds.

## Materials and Methods

### Strains and Growth conditions

*Aggregatibacter actinomycetemcomitans* ATCC 43718 and *Streptococcus mutans* ATCC 25175 were used in all described experiments. The strains were maintained on blood agar No2 (Oxoid, Hampshire, UK) supplemented with hemin (5 mg/mL) (Sigma Aldrich, Belgium), menadione (1 mg/mL) (Sigma Aldrich, Belgium) and 5% sterile horse blood (Oxoid, Hampshire, UK) or cultured in liquid medium in Brain Heart Infusion (BHI) (Carl Roth, Belgium) broth under aerobic conditions at 37°C.

### Flow cytometric measurements

#### Sample preparation and treatment

After overnight culture in the conditions that were previously described, the bacterial cultures were measured by flow cytometry and SYBR Green/ Propidium Iodide (SGPI) staining to determine the intact cell concentration. In more detail, the samples were diluted in sterile (and filtered through 0.2 um) PBS (PBS tablet, Merck, Belgium) and stained with the nucleic acid stain SYBR^®^ Green I and Propidium Iodide that stains permeabilized cells^30^. SYBR Green I (10,000X concentrate in DMSO, Invitrogen) was diluted 100 times in 0.22 μm-filtered DMSO (IC Millex, Merck, USA) and Propidium Iodide (20 mM in dimethyl sulfoxide (DMSO), Invitrogen, USA) was diluted 50 times. Samples were stained with 10 μL/mL staining solution. Next, they were incubated at 37°C for 20 min and measured with a benchtop Accuri C6+ cytometer (BD Biosciences, Belgium).

Overnight cultures were subsequently diluted in sterile dH2O (for *S. mutans*) or sterile Evian bottled water (Evian, France) (for *A. actinomycetemcomitans*) to a final concentration ~1×10^6^ cells/mL according to the previous measurement and further stained with SGPI with the above mentioned approach. Just before measurement the corresponding concentration of chlorhexidine digluconate (Merck, Belgium), cetylpyridinium chloride (Carl Roth, Germany) or triclosan (Merck, Belgium) was added to the diluted culture and mixed well.*A. actinomycetemcomitans* was subjected to CHX or CPC in 2 mg/mL, 10 mg/mL or 20 mg/mL. *S. mutans* was subjected to CHX, CPC and triclosan either in 2 mg/mL or in 20 mg/mL.

#### Sample measurement/ instrument settings

All samples were measured with a benchtop Accuri C6+ cytometer (BD Biosciences, Belgium). The stability of the instrument was verified daily using CS&T RUO beads (BD Biosciences, Belgium). The blue laser (488 nm) was used for the excitation of the stains. The filters for the (fixed gain) photomultiplier detectors used during the measurements were 533 nm with a bandpass of 30 nm for the green fluorescence (FL-1) and 670 nm longpass filter for the red fluorescence (FL-3). The threshold was set on the 533/30 nm (FL-1) detector at the arbitrary unit of 1200 and on the 670 nm(FL-3) at the arbitrary unit of 1200. The threshold was decided based on measurements of control samples (growing culture, heat-killed culture, medium and dH2O/diluent) in order to avoid background noise and allow for maximum measurement of total events in the same acquisition. Sample acquisition took place at a flow rate of 66 μL/min continuously time of 15 min.

### Data analysis

#### Data filtering and cleaning

Flow cytometric data were analyzed in R (version 4.0.3). The Phenoflow’s (v1.1.2)^31^ function *‘time_discretization’* was used to concatenate the files into smaller files of fixed time frames of 30 seconds.

Two different gating strategies were used for each species to gate a. intact cells, b. damaged cells and c. total cells (Supplementary Figure 1 & 2). Non-treated and heat-killed samples were used as control samples to define the intact and damaged cell gates.

#### Time and Dose response Analysis

Intact cell numbers, based on the ‘intact cells’ gate, were used for calculating the killing rate under the different antimicrobials. First the percentage of surviving cells was calculated as the ratio of intact cells at each time point over the average intact cells of non-treated control samples. Subsequently, the *‘drm’* function from the drc package (v3.0.1)^32^ was used to fit different log-logistic models to the time-effect or dose response data. For all log logistic models the maximum asymptote was constraint to 100. The fit of the different models was evaluated with the *‘mselect* function of the drc package to identify the model that best fitted the data based on the Akaike’s information criterion (AIC) and the lack-of-fit test (against a one-way ANOVA model). The model with the lowest AIC and highest p-value in the lack-of-fit test was chosen. The Hill coefficient and the effective time 50 or effective dose 50 were calculated from the model. Effective time/dose 50 is the time or dose for which 50 per cent of killing is reached.

#### Phenotypic sub-populations PhenoGMM

The *‘PhenoGMM’* function^24^ of the PhenoFlow package was used to determine phenotypes to which cells can be assigned. Initially, the background was removed by applying the ‘total cells’ gate (Supplementary Figure 2). Subsequently the ‘PhenoGMM’ function was applied on a representative subset of the data, using the FSC-H, SSC-H, FL1-H, and FL3-H parameters as input. The fcs samples were resampled with replacement to 1,000 events. The best number of mixtures/phenotypes to describe the dataset was chosen from 1:20 based on Bayesian Information Criterion (BIC). Increasing the number of allowed phenotypes was leading to a higher number by BIC but eventually 20 phenotypes limit was chosen as a number that allowed to capture the data variation without increasing the complexity and result interpretation. After the Gaussian Mixture Model wasfitted, it was used to calculate the abundance of each mixture in the total dataset.

## Data Availability

Flow cytometry data (.fcs format) are available on the FlowRepository archive under repository ID FR-FCM-Z4VR.

## Acknowledgments

This work was supported by ‘Fonds voor Wetenschappelijk Onderzoek’—FWO (G0B2719N & 3G020119). R.P. is supported by a postdoctoral fellowship of the ‘Fonds voor Wetenschappelijk Onderzoek’ —FWO (project 1221020N).

## Supplementary material

Real-Time Flow Cytometry to assess qualitative and quantitative responses of oral pathobionts during exposure to antiseptics

### Supplementary Figures

**Figure S1:**
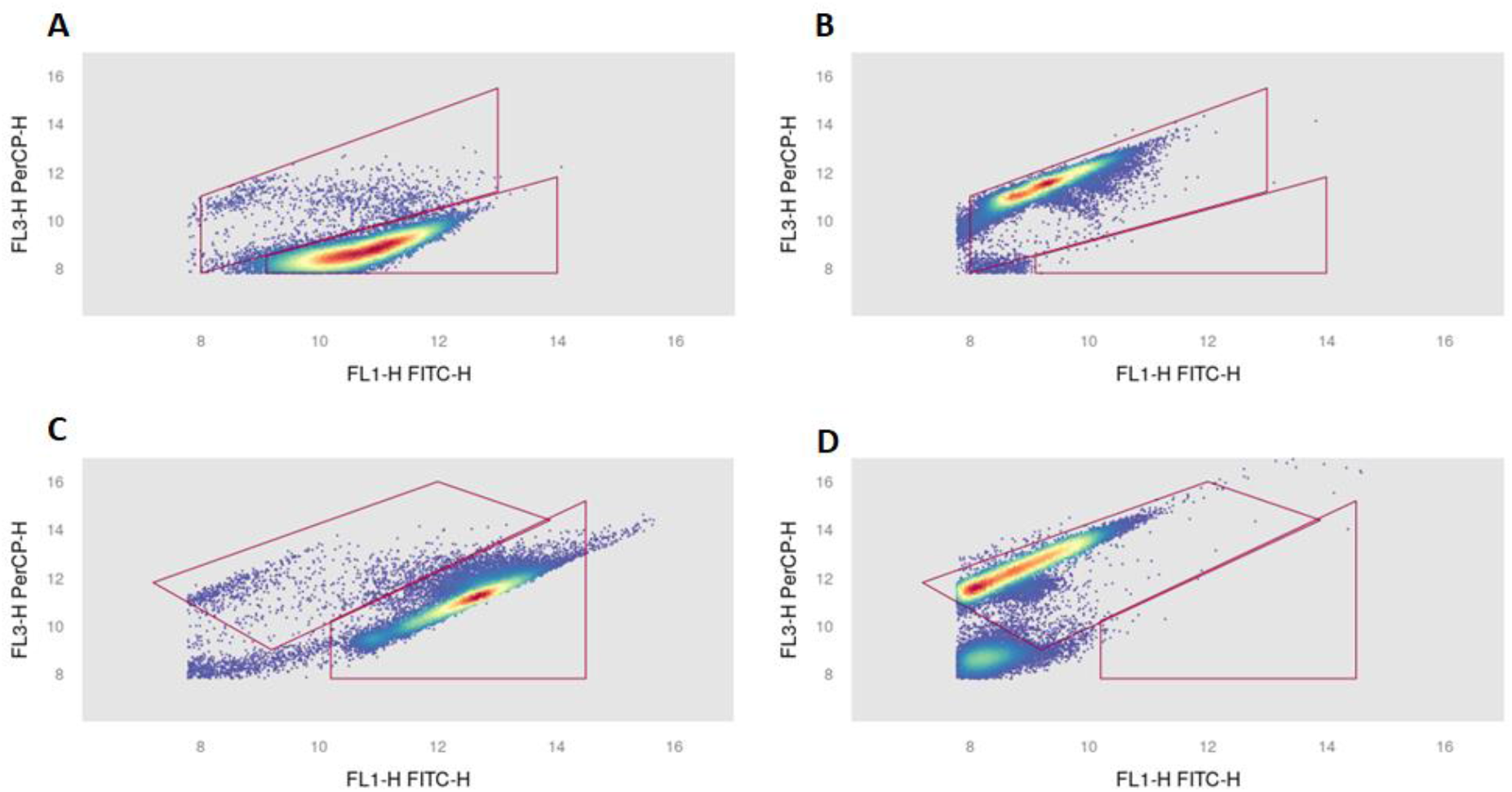
Examples of the manual gates that were drawn in the FL1/FL3 dot plots for the intact and damaged cell populations based on non-treated control (A, C) and heat-killed cells (B, D) for *A. actinomycetemcomitans* (A-B) and *S. mutans* (C-D)

**Figure S2:**
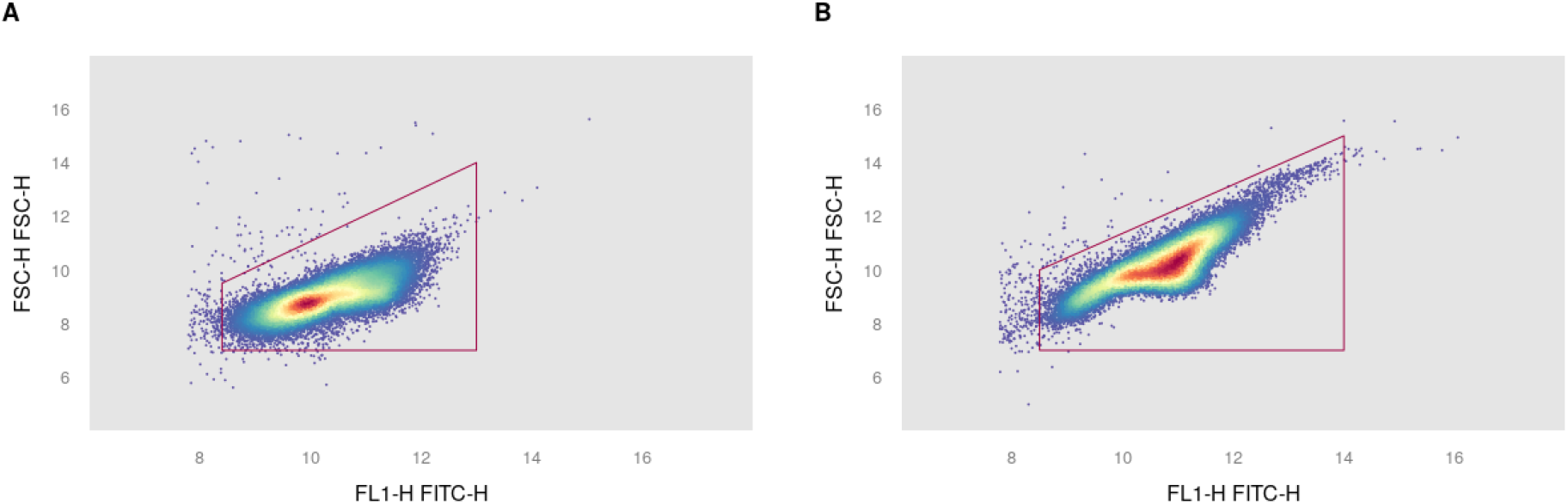
Examples of the manual gates that were drawn in the FL1/FSC dot plots for total cell populations to distinguish cells from background for *A. actinomycetemcomitans* (A) and *S. mutans* (B)

**Figure S3:**
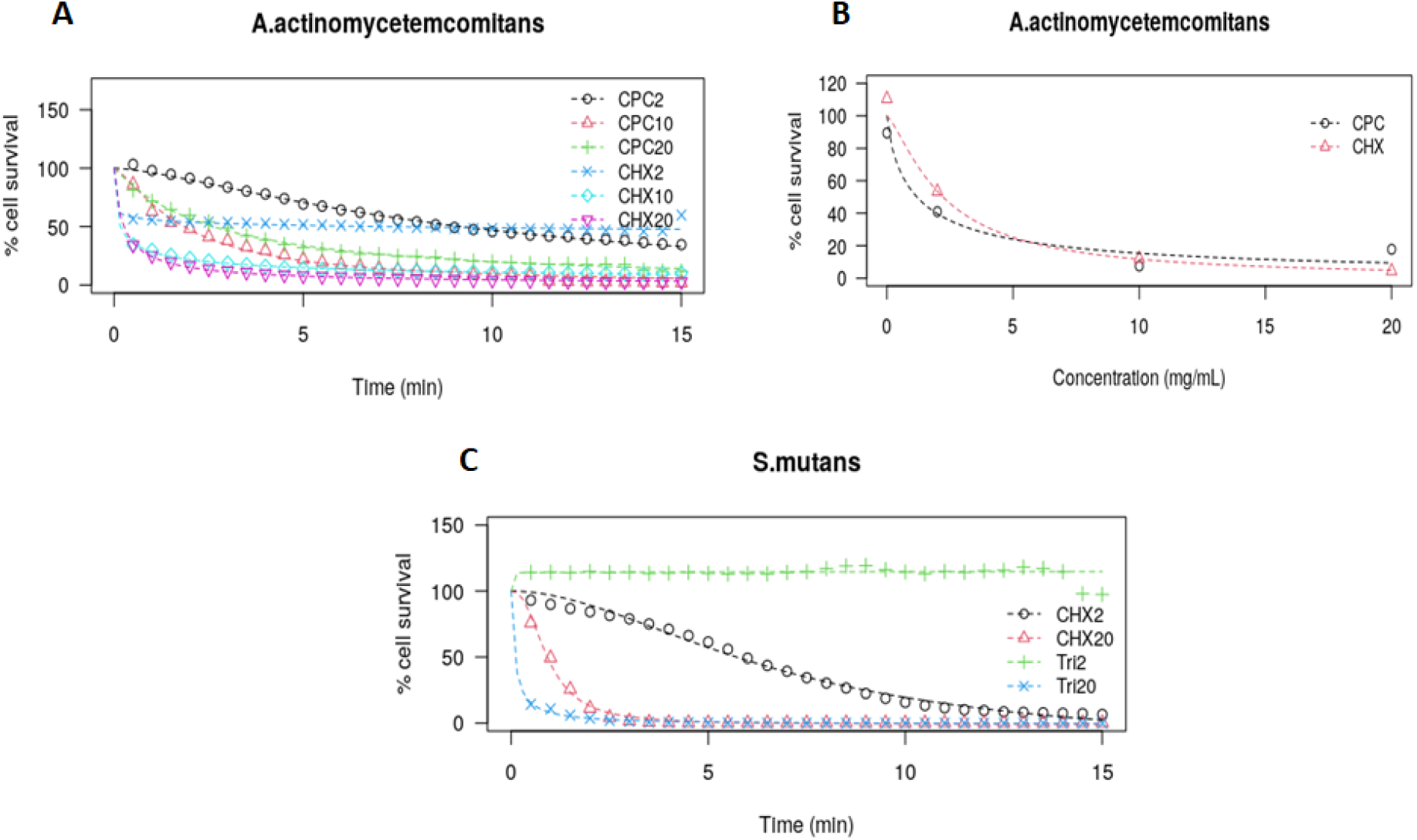
The log-logistic models fit on the percentage of surviving cells for a) *A. actinomycetemcomitans* over time for the different treatments (3 parameter log-logistic model) b) *A. actinomycetemcomitans* in different concentrations of the antiseptic at 10 min of treatment (3 parameter log-logistic model) and c) *S. mutans* over time for the different treatments (4 parameter log-logistic model)

**Figure S4:**
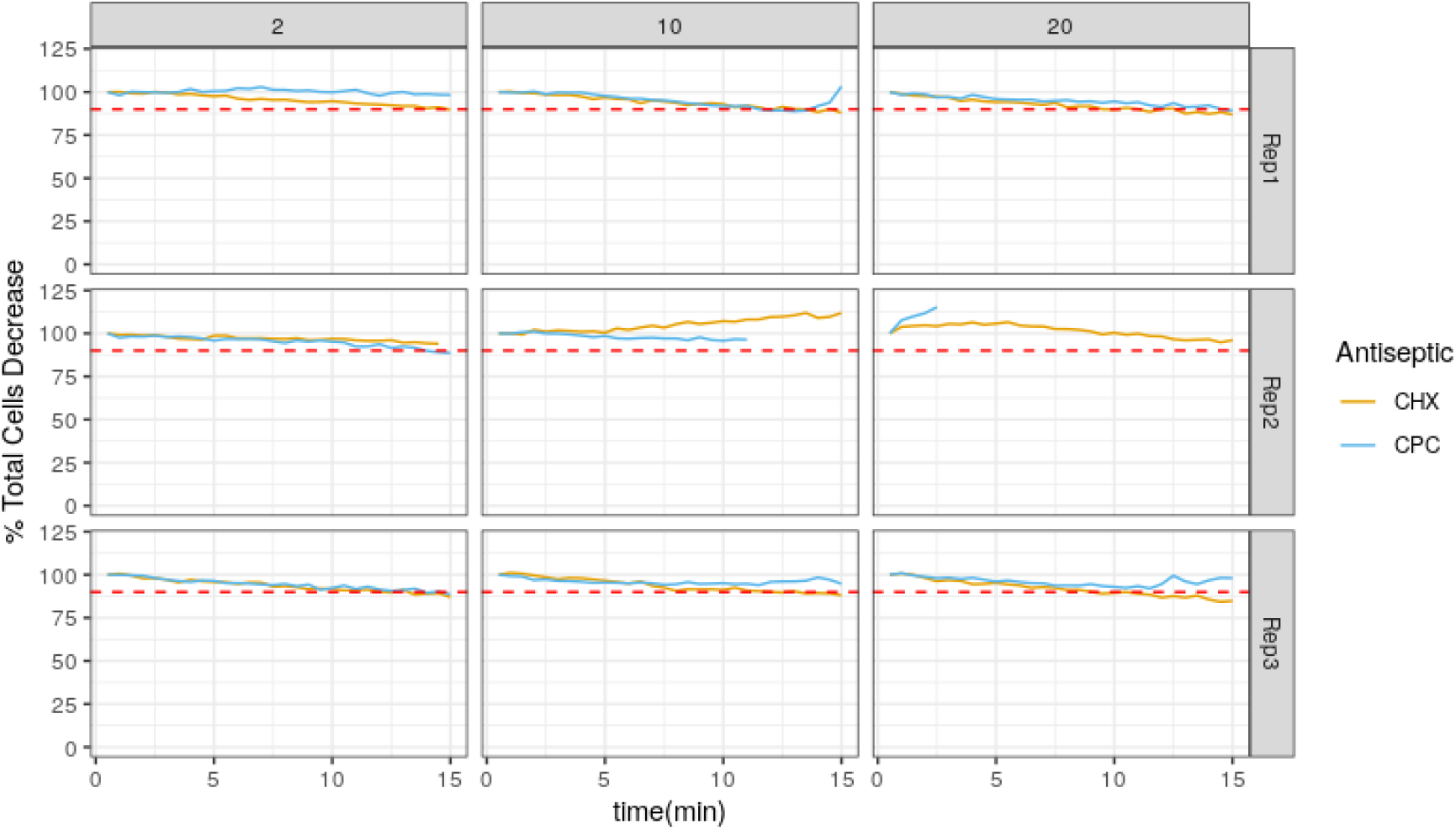
The percentage of total cells (intact and damaged cells) of *A. actinomycetemcomitans* over time compared to the first time point, for the three antiseptic concentrations and three replicates. The red dotted line represents 90% of the initial total cells.

**Figure S5:**
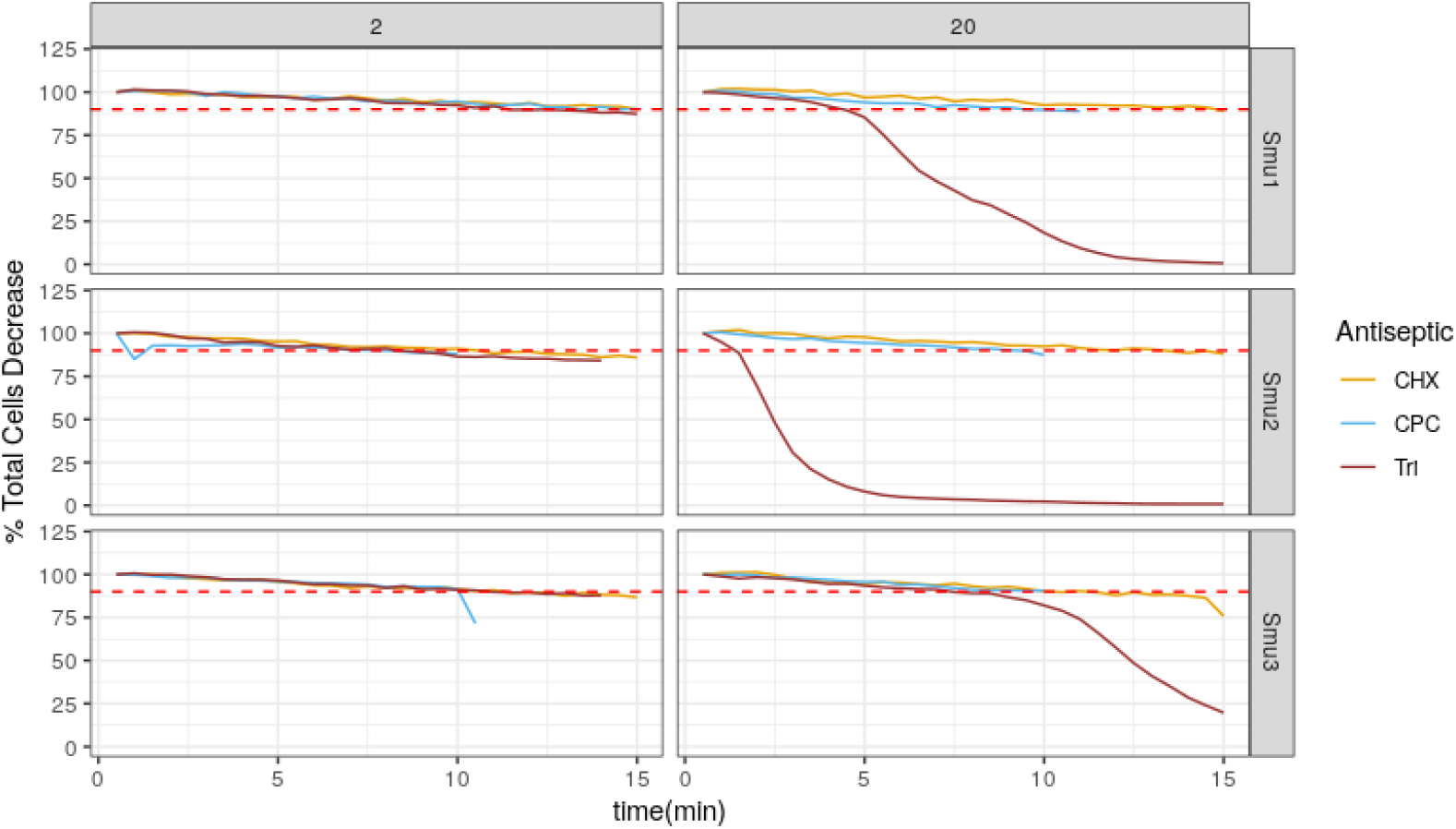
The percentage of total cells (intact and damaged cells) of *S. mutans* over time compared to the first time point, for the two antiseptic concentrations and three replicates. The red dotted line represents 90% of the initial total cells.

**Figure S6:**
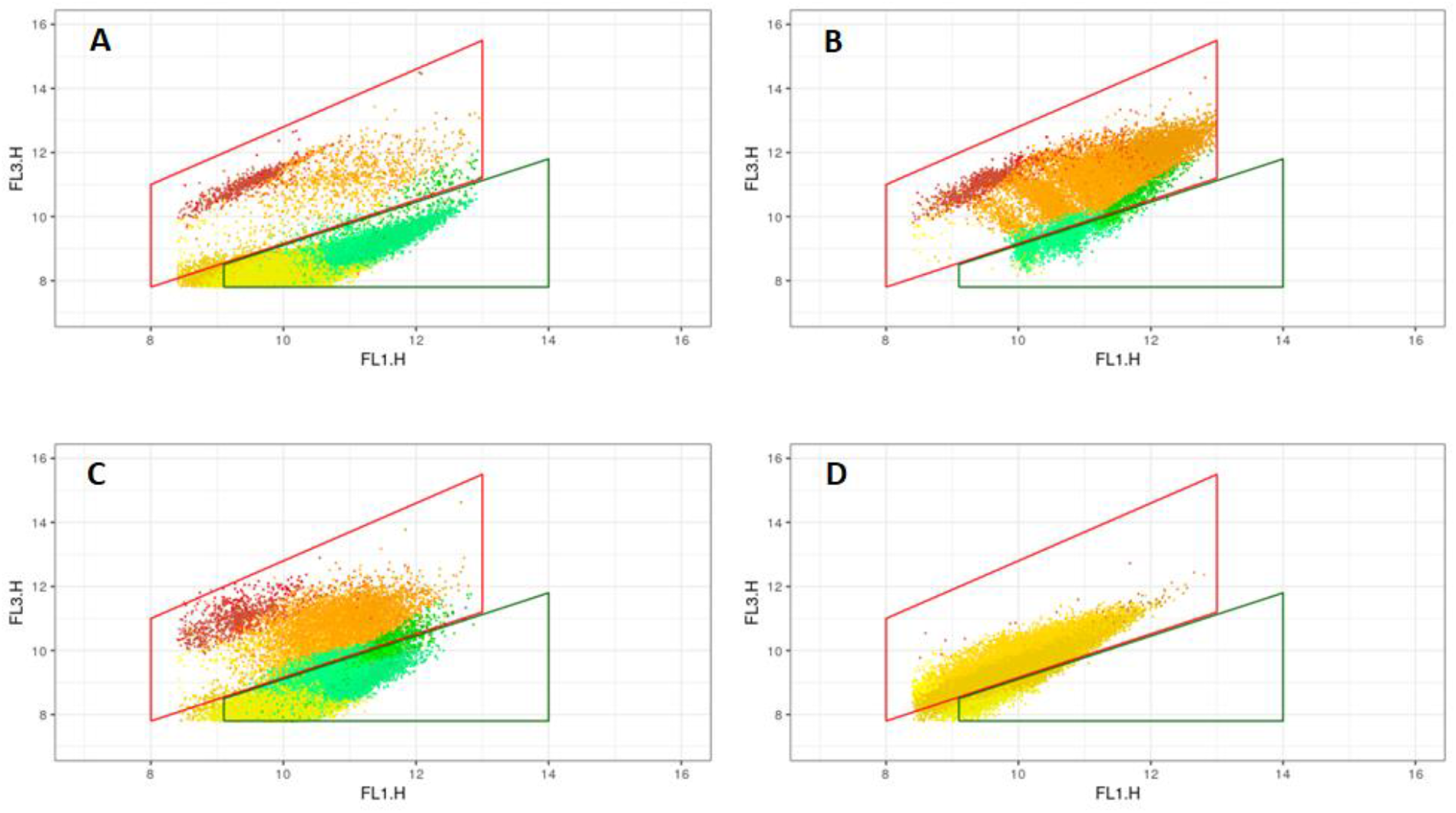
Examples of *A. actinomycetemcomitans* cells coloured according to the phenotypes as estimated based on the Gaussian Mixture model: a) untreated control after 4 min, B) cells treated with 20 mg/mL CHX after 4 min, C) cells treated with 2 mg/mL CPC after 4 min and D) cells treated with 20 mg/mL CPC after 4 min. The gates represent the manual gates that were drawn for the first part of the experiment: a) green for the intact cells and b) red for the damaged cells.

**Figure S7:**
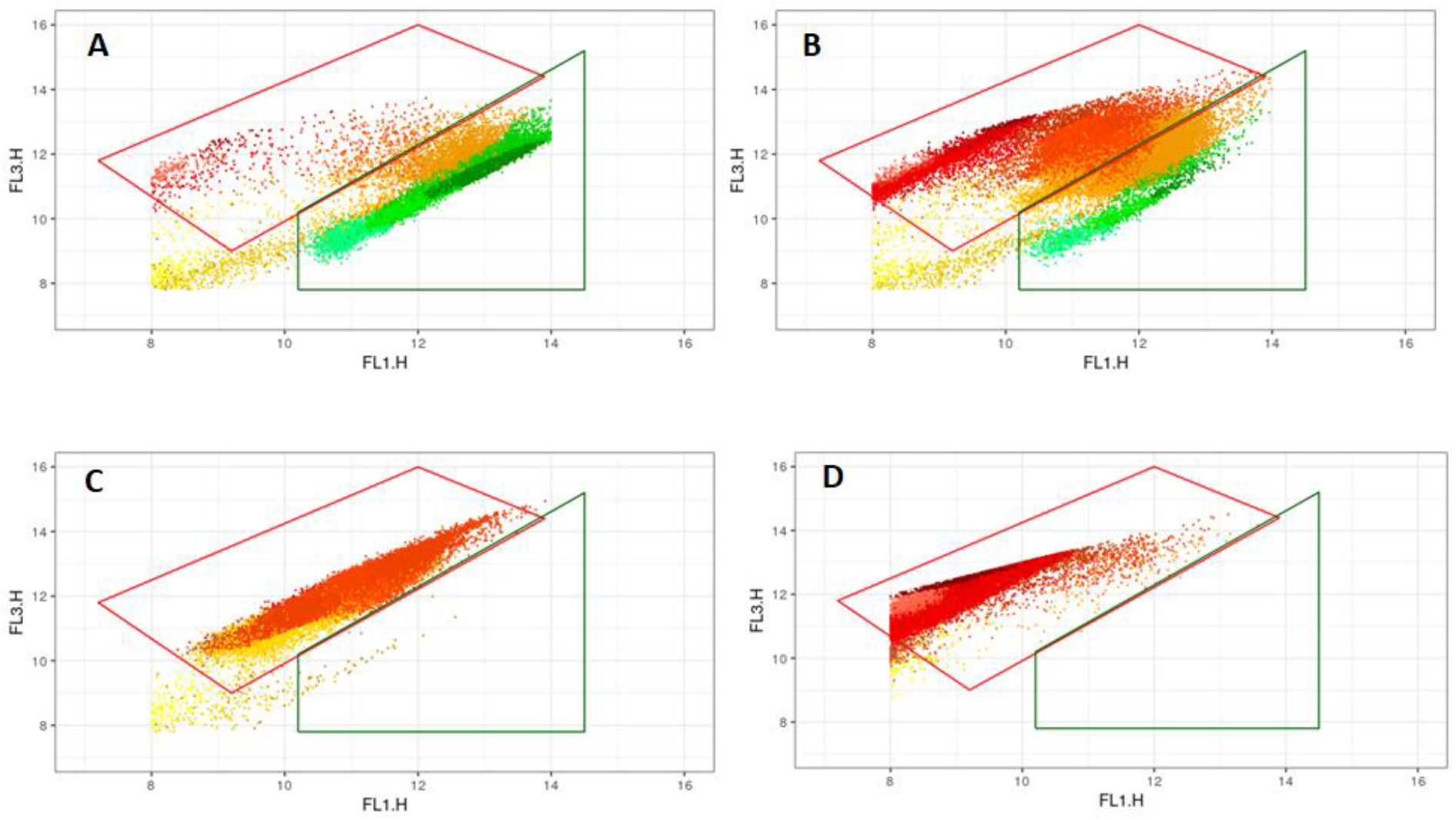
Examples of *S. mutans* cells coloured according to the phenotypes as estimated based on the Gaussian Mixture model: a) untreated control after 4 min, B) cells treated with 2 mg/mL CHX after 4 min, C) cells treated with 20 mg/mL CPC after 4 min and D) cells treated with 20 mg/mL triclosan after 4 min. The gates represent the manual gates that were drawn for the first part of the experiment: a) green for the intact cells and b) red for the damaged cells.

